# Characterization of light penetration through brain tissue, for optogenetic stimulation

**DOI:** 10.1101/2021.04.08.438932

**Authors:** Emily L. Johnson, Darren Walsh, Frances Hutchings, Rolando Berlinguer Palmini, Nikhil Ponon, Anthony O’Neill, Andrew Jackson, Patrick Degenaar, Andrew J. Trevelyan

**Affiliations:** Institute of Biosciences, Medical School, Framlington Place, Newcastle University, Newcastle upon Tyne, NE2 4HH, UK; School of Computing, Newcastle Helix, Newcastle University, Newcastle upon Tyne, NE4 5TG, UK; School of Engineering, Merz Court, Newcastle University, Newcastle upon Tyne, NE1 7RU, UK

## Abstract

The recent development of optogenetic tools, to manipulate neuronal activity using light, provides opportunities for novel brain-machine interface (BMI) control systems for treating neurological conditions. An issue of critical importance, therefore, is how well light penetrates through brain tissue. We took two different approaches to estimate light penetration through rodent brain tissue. The first employed so-called “nucleated patches” from cells expressing the light-activated membrane channel, channelrhodopsin (ChR2). By recording light-activated currents, we used these nucleated patches as extremely sensitive, microscopic, biological light-meters, to measure light penetration through 300-700µm thick slices of rodent neocortical tissue. The nucleated patch method indicates that the effective illumination drops off with increasing tissue thickness, corresponding to a space constant of 317µm (95% confidence interval between 248-441µm). We compared this with measurements taken from directly visualizing the illumination of brain tissue, orthogonal to the direction of the light. This yielded a contour map of reduced illumination with distance, which along the direction of light delivery, had a space constant, *τ* 453µm. This yields a lower extinction coefficient, µ_e_ (the reciprocal of *τ*, ∼3mm^-1^) than previous estimates, implying better light penetration from LED sources than these earlier studies suggest.

## Introduction

The development and design of optogenetic brain machine interfaces (Deisseroth, 2011; Paz et al., 2013; Zaaimi et al., 2021), for clinical use, will rely heavily on accurate models of the different biological, physical and engineering elements. One critical issue is how well light penetrates through brain tissue to activate the opsins, because this will dictate the extent of influence of an implanted light source on the network. Light penetration is affected by reflection and refraction at the point of entry into the tissue, and subsequently by scattering and absorption effects. The Beer-Lambert law states that the radiant energy will decay with 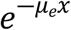, where *μ*_*e*_ is the extinction coefficient and x is the distance traversed through the tissue. The extinction coefficient *μ*_*e*_ is the sum of the scattering *μ*_*s*_ and absorption *μ*_*a*_ coefficients. It is important to measure the coefficient value specific to the wavelength of the relevant opsin, and for the brain tissue in question, and also take into consideration the type of light source. Previous attempts to characterize the optical properties of tissue have mostly used collimated light sources (Aravanis et al., 2007; Dong et al., 2018; Yona et al., 2016). However, implantable light emitters will have beam profiles which deviate significantly from collimated beams. A further issue is that the measurements in these experiments typically utilized macroscopic light sensors, integrating almost instantaneously (Aravanis et al., 2007; Yona et al., 2016). In optogenetic applications, though, the effective light sensor could either be considered to be the opsin molecule, or alternatively the cell, both of which are microscopic, and integrate light over milliseconds. Both the size and integration properties of the sensors may affect the estimates of how well light penetrates through the tissue. We were interested, therefore, to revisit this question of light penetration through interposed tissue, employing opsin molecules as sensors – that is to say, using the actual biosensors that would be used in any actual implementation of optogenetic brain control.

Opsins are well suited for this purpose: they constitute highly sensitive light detectors, which transduce the light into a readily recordable electrical signal. The task, however, is more difficult than simply recording a neuron at a set distance from the light source, because neurons have extensive processes, which represent a very distributed physical structure in the tissue, with the different elements at variable distance from the light source. These different neuronal processes also vary greatly in their excitability, and the subsequent transduced effect on the membrane potential is then filtered and summated by the electronic structure of the cell; consequently, the recorded signal is a complex function of the direct effect of the light on the optogenetic proteins.

To measure light penetration, ideally, one would like to use a point light-sensor, which is clearly not the case for an intact neuron. Nor can this be achieved easily using outside-out patches, since the number of opsin proteins in an outside-out patch is typically tiny, meaning that the response is quantal, and ill-suited for mapping out a light-response curve. A different technique, in which the entire neuronal soma is pulled off with a patch electrode – termed a “nucleated patch” (Bekkers, 2000; Gurkiewicz and Korngreen, 2006) – creates a minuscule structure, typically around 10µm across, but which may have hundreds to thousands of opsin molecules within the membrane. These recordings can, in principle, generate both small and large light-induced currents, depending on the level of illumination, but it is necessary still to calibrate these responses, since each nucleated patch differs in its content of opsin molecules. To do this, we created a recording chamber in which a nucleated patch could be illuminated from two different sources: for one of these, the microscope, we were able to measure accurately the illumination power onto the nucleated patch, and this provided the calibration for each recording; light from the other source, an LED, passed through brain slices of different thicknesses to illuminate the nucleated patch. We measured light-response curves for both light sources, for each nucleated patch, and calculated the discrepancy (the “correction”) between the two light responses, to derive the progressive reduction in light penetration for increasing distance travelled through brain tissue. We compare these results with measures taken from directly visualizing scattered light, orthogonal to the direction of illumination (Dong et al., 2018).

## Methods

### Nucleated patch-clamp recordings

All animal handling and experimentation was done according to UK Home Office guidelines and the requirements of the United Kingdom Animals (Scientific Procedures) Act 1986. Mice were housed under a 12:12 h light/dark cycle with free access to food and water. All efforts were made to minimize animal suffering and the number of animals used.

#### Dissociated neuronal cultures

All cells were maintained in a humidified incubator at 37°C, CO_2_ (5%): air (95%), and all culture reagents were purchased from Gibco, (ThermoFisher Scientific, Paisley, UK), unless otherwise stated. Primary dissociated neuronal cultures were prepared from embryonic rat pups (E18-20). Pregnant Sprague–Dawley dams (<350g) were sacrificed by cervical dislocation and pups were removed and decapitated. Neocortical and hippocampal tissue was dissected and transferred to ice cold dissection media (DM, HBSS + Ca^2+^/Mg^2+^, 100mM HEPES, 1mM glucose). For enzymatic dissociation, tissue was incubated with papain (2units/ml, diluted in DM, Sigma Aldrich) at 37°C for 40mins, washed with DM and then culture medium (Neurobasal A, 2% B-27 supplement, 0.5% foetal bovine serum (FBS), 0.5% glutamate, 0.5% antibiotic–antimycotic). Cells were suspended in culture medium (10ml) and manually triturated using serological pipettes of decreasing diameters (25ml, 10ml, 5ml). Cells were counted using a haemocytometer, diluted and plated (1×10^5^ cells in 0.5ml culture medium per well) into 24 well plates containing sterile glass coverslips (thickness ∼0.19mm; Thermo Scientific) pre-coated overnight with poly-D-lysine (10µg/ml diluted in ddH_2_0, Sigma-Aldrich, Gillingham, UK). Culture medium was replaced 24hrs post-plating and subsequently half-changed every 3 days *in vitro* (DIV), with culture medium lacking FBS to minimise glial cell proliferation.

To achieve ChR2 expression, at DIV 7, cells were incubated with a 3^rd^ generation lentiviral vector (LVV) encoding ChR2 (hSyn-EYFP-ChR2(H134R), ABM Inc, Vancouver, Canada).

The LVV was diluted in culture medium to a final concentration of 10 MOI (i.e. 10 viral particles per cell). Cells were used for experiments between DIV 12-17.

#### Acute brain slice preparation

Adult C57/BL6 mice (2-4 months) were sacrificed by cervical dislocation and brains were immediately transferred to ice-cold oxygenated (95% O2/5% CO2) artificial cerebrospinal fluid (ACSF: 125mM NaCl, 26mM NaHCO_3_, 10mM glucose, 3.5mM KCl, 1.26mM NaH_2_PO_4_, 3mM MgCl_2_). Coronal brain slices (300, 500 and 700µm thickness) were prepared using a vibrating microtome (LEICA VT1200; Leica Microsystems (UK) Ltd, Milton Keynes, UK). Slices were then transferred to a oxygenated submerged incubation chamber containing ACSF (as above except with 1mM MgCl_2_ and 2mM CaCl_2_).

#### Patch-clamp electrophysiology

Segments of coverslips with cultured neurons transduced with LVV-hSyn-EYFP-ChR2(H134R) were transferred to the recording chamber, alongside acute brain slices fixed by a harp directly over the CREE LED. Cultured neurons expressing the EYFP fluorescent marker were selected under direct visual guidance (SLICESCOPE, Scientifica, Sussex, UK) fitted with a CooLED pE epifluorescence imaging system (Andover, UK) and recorded in whole-cell patch-clamp mode. For all recordings, cells and brain slices were continuously perfused with oxygenated ACSF (32°C). Patch-pipettes with a resistance of 5-7 MΩ were made from borosilicate glass capillary tubes (0.86mm internal diameter; Harvard Apparatus, Cambridge, UK) using an electrode puller (P-87; Sutter Instrument Co, CA, USA). Patch-pipettes were filled with intracellular solution (K-methyl-SO_4_ 125mM, Hepes 10mM, Mg-ATP 2.5mM, NaCl 6mM). Both the microscope objective and headstage positioning were controlled by individual micromanipulators (Patch star PS-700C; Scientifica, East Sussex, UK) enabling precise movements over three axes (x, y and z). Patch-clamp recordings were made using an Axopatch 700B amplifier/Digidata 1440A interface (Axon Instruments; Foster City, CA, USA), controlled by Clampex 10.5 software (Molecular Devices; Foster City, CA, USA). Signals were sampled at 10 kHz and low-pass filtered at 2 kHz.

#### Nucleated patch recordings

after achieving whole-cell recording mode and verifying that the cell expressed ChR2 through a test illumination, a “nucleated patch” was pulled (Figure 1C) (Bekkers, 2000; Gurkiewicz and Korngreen, 2006). In brief, this was done by having a very gentle suction (∼<20psi), and drawing the electrode directly back, very slowly (over ∼1-2mins), until the nucleated somatic bleb, on the end of the electrode, separates from the rest of the cell. This typically happens when the electrode tip has been drawn back about 50-100µm. This technique creates a large, outside-out patch, containing the nucleus of the cell, which displays macroscopic currents, and, importantly, also allows one to move the patch electrode. The nucleated patch could thus be relocated precisely above the CREE-LED, and separated by neural tissue of a specified width.

**Figure 1.**
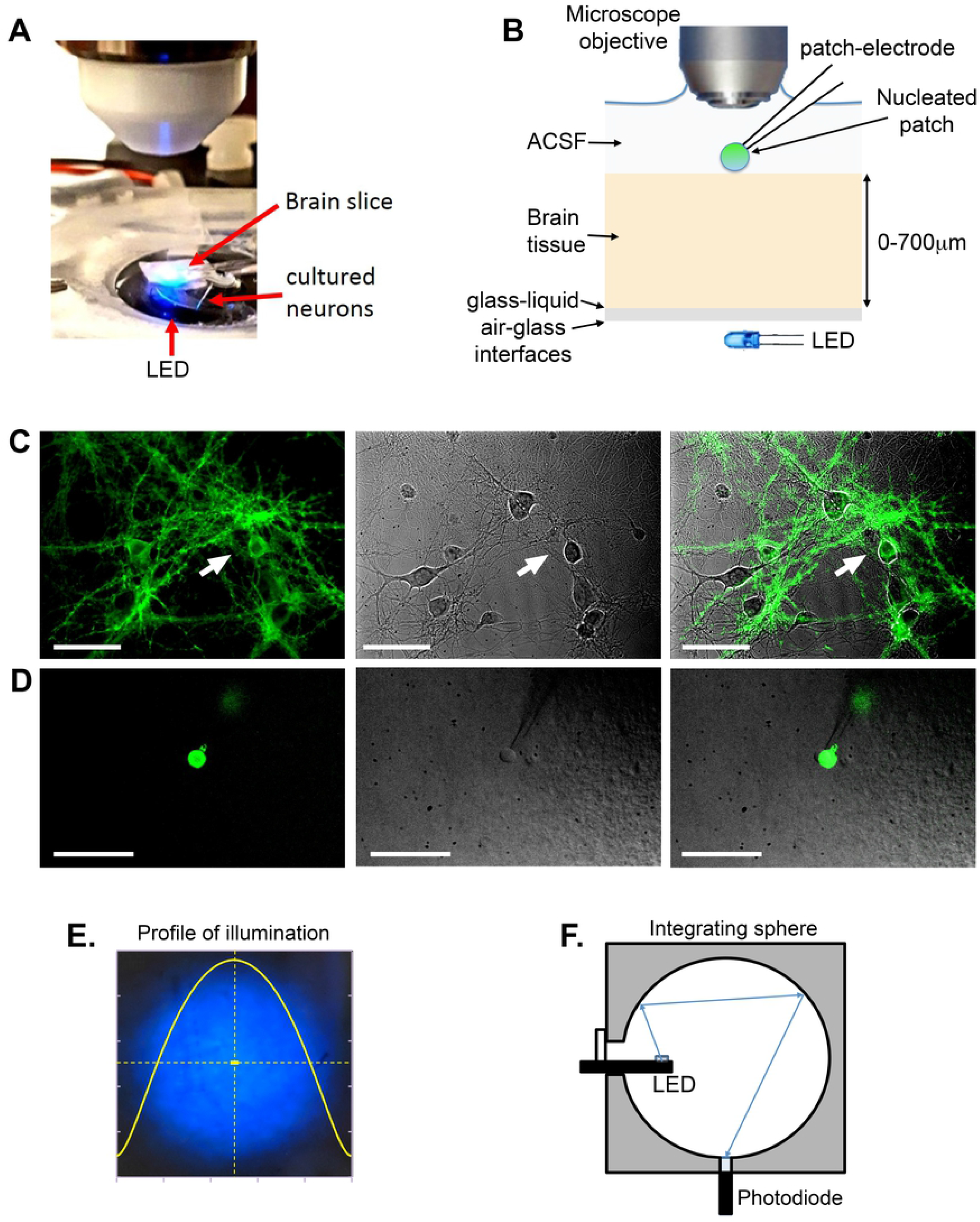
Using nucleated patch recordings to measure optogenetic activation through brain tissue. (A) Photograph of the recording arrangement, showing a glass cover slip of cultured neurons expressing ChR2 in the recording chamber, together with a brain slice of specified thickness (300 / 500 / 700µm). The LED was placed below glass at the bottom of the recording chamber, and also underneath the brain slice, allowing light to be delivered either from below, from the mini-LED, or above, through the microscope objective. (B) Schematic of the recording arrangement, as viewed from the side. (C) Cultured neurons expressing ChR2 tagged to eGFP (left panel, epifluorescence; middle panel, DIC; right panel, combined images). (D) Same visualization of a nucleated patch, pulled from one of the labelled cultured neurons, and then positioned just above the brain slice. (E) Profile of illumination delivered through the microscope objective. (F) Measuring the mini-LED light intensity with an integrating sphere.

#### Illumination sources

The mini-LED illumination was provided by a CREE DA2432 LED (470 nm emission peak, dimensions: 320 × 240 µm). The mini-LED was mounted on a silicon shaft and sealed using a Poly-DiMethyl-Siloxane (PDMS) silicone sealant as per (Dong et al., 2018). The probe was placed below the recording chamber, which was itself mounted on a light microscope (BX61, Olympus), as shown in Figure 1A,B. The mini-LED was powered by an isolated pulse stimulator (2100, A-M SYSTEMS, Hinckley, UK); enabling control of the amplitude, frequency and duration of the current. Brain slices of different thickness were placed above the mini-LED, as illustrated in Figure 1A. This configuration enabled illumination of the tissue from underneath using the mini-LED or from above using the microscope LED illumination (465nm). The LED light intensity at its source was estimated using a 50mm integrating sphere (Laser2000) configuration (Figure 1F). A conversion of 1.05 mW/mA was achieved in the range 1-5mA with minimal droop effects. Though This was to provide a ballpark figure of intensity, as the final calculation of the space constant for light penetration through the tissue was performed by normalizing the mini-LED illumination. This normalization was done in part, also to address factors such as the reflectance at several interfaces (air/glass/water – see the schematic diagram in Figure 1B) that lay between the mini-LED and the nucleated patch, and which were difficult to quantify experimentally, but which were constant for all experiments. The mini-LED intensity was altered over a 5-fold range, for the nucleated patch illumination.

The microscope illumination was provided by a CooLED epifluorescence system (Andover, UK), allowing the illumination to be varied over a 100-fold range. The illumination was further modulated by placing neutral density filters (Thorlabs; 64-fold reduction) within the microscope light path (Total light range ∼6400-fold).

#### Light intensity at focal point

We first measured the total power of the epifluorescence light delivered through the 40x PlanFluor microscope objective, without neutral density filters, using a Thorlabs PM120VA photodiode sensor. The spread of light at the focal plane was photographed, and the cross-section was approximated to a Gaussian distribution (Figure 1E). The light intensity onto the nucleated patch was then estimated to be the fraction of the total power (the integral from –∞ to +∞), subsumed by the central 10µm (∼diameter of a nucleated patch pulled from a putative pyramidal cell) (∼0.1% of the total power). Peak irradiance was estimated to be 26.4 mW/mm^2^, at 100% microscope illumination power and without any neutral density filters. The nucleated patch could be moved freely, since the electrodes were mounted on Patchstar micromanipulators (Scientifica Ltd); for all light responses, we moved the nucleated patch to the centre of LED illumination (which could be estimated very accurately through the microscope), and since it was also centred within the field of view of the microscope, this effectively aligned the peak illumination from both sources.

#### Data analysis

traces were down-sampled by factor of 10, low pass filtered using a -3.0dB cutoff at 40Hz. Mean and peak photocurrents over the 200ms illumination period were calculated, from 2-5 repeat trials per nucleated patch.

### Light penetration experiment

Figure 4 (A-C) shows the light penetration measurement experimental setup. Brains were extracted from 6 weeks old C57BL6 mice, after cervical dislocation. The brains were then dissected in two hemispheres along the sagittal line. Each hemisphere was subsequently cut along the dorsal surface forming a 90-degree angle between the dorsal and sagittal planes. The brain hemisphere was placed in a microscope (Olympus BX 61, upright) stage chamber submerged in ACSF (Figure 4c) at room temperature with the dorsal surface (b) facing upwards for observation through the microscope lens (Olympus 2.5X NA, (f). The sagittal plane (a) was placed facing a 400um diameter cannula tip (d) connected to a LED via a fibre optic (e). Illumination was performed using CoolLEDs precise Excite LEDs (470 nm, 585 nm). Light penetration measurements were taken with an Andor iXon DV887 back illuminated EMCCD camera and the images of the brain dorsal surface acquired with Andor Solis software.

## Results

### Nucleated patch biosensor method for measuring light penetration

We took two different approaches to measuring light penetration through blocks of mouse brain tissue. The first, used cultures of neurons expressing the light-activated opsin, ChannelRhodopsin-2 (ChR2) to create microscopic biosensors for measuring light penetration through different thicknesses of brain tissue. The experimental configuration is illustrated in Fig 1. Nucleated patches were pulled from cultured neurons, expressing ChR2. The electrode, with the attached nucleated patch, was then moved away from the cultures, to another location directly above an LED light source that was mounted below the recording chamber. In most recordings, there was additionally a slice of mouse brain tissue between the light source and the nucleated patch, ranging between 300-700µm thick. Other recordings were also made without brain slices (“0µm thickness”, although note that the distance from the LED source was approximately the same as for recordings with 300µm thick brain slices), to allow other causes of reduced illumination to be estimated. The nucleated patch was positioned at the focal point for the microscope objective, allowing us also to illuminate it from above, in a highly controlled fashion, using the epifluorescence light path of the microscope. Importantly, this meant that the LED and the microscope illuminations were both aligned, and centred on the nucleated patch, since this could be visualised, and translocated precisely by the micromanipulators controlling the electrode.

We delivered 250ms steady-state light illumination (square pulses of light delivery), at different intensities, generating reliable light responses with a large amplitude peak current occurring within the first 50ms, which then desensitised (Fig 2A). At the higher currents surface temperatures on the LEDs can rise several degrees over this timescale in air. But as it was separated from the tissue medium, we have assumed no heating effects. Similarly, no heating effect was expected from the microscope illumination. The expected optical irradiance was significantly below the threshold for optically induced heating seen by Stujenske (Stujenske et al., 2015).

**Figure 2.**
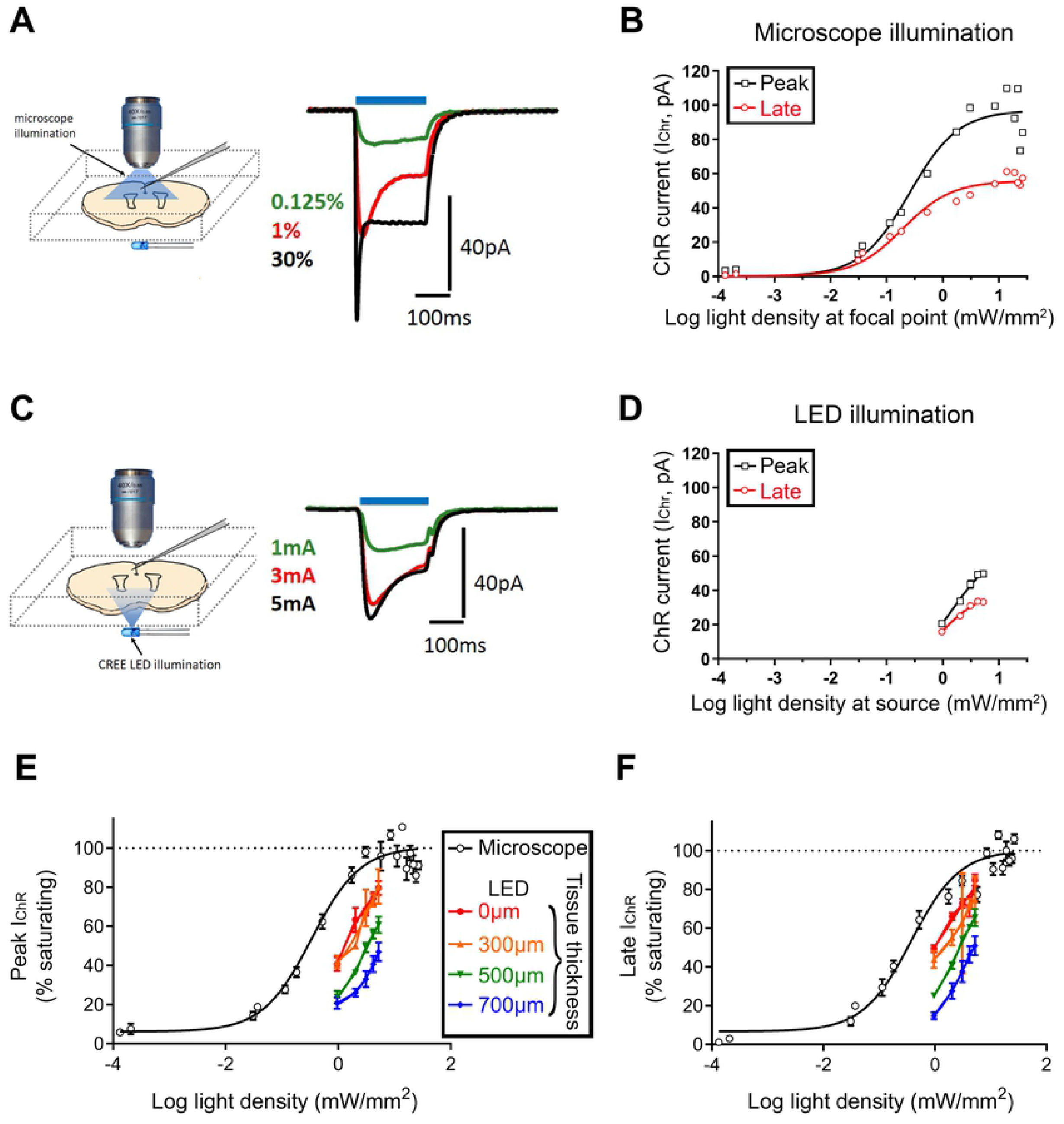
Microscope and LED illumination induce equivalent light-response curves. (A) Three different light responses, from a nucleated patch clamp, induced by optically activating ChR2 from above, through the microscope objective (units represent the %age of maximal illumination (no neutral density (ND) filters). Maximal illumination, at the focal point, where the nucleated patch was located, was measured to be 26mW/mm^2^. (B) The light response curve, for the same nucleated patch, measured either for the peak current, or for the mean of the final 100ms of illumination (an approximation of the steady state current). (C) Example light responses from the same nucleated patch, when illuminated from below, through 700µm of brain tissue. The units represent the different currents passed through the LED. (D) The light response curve for the LED illumination for this nucleated patch, but plotted relative to the measured output of the LED at its source. The discrepancy between these measures, and those made using the microscope illumination (B), is because the light from the LED additionally passes through air, glass and brain tissue (see Fig 1B). (E) Pooled data from 4 nucleated patches, for the peak currents. (F) Pooled data from 4 nucleated patches, for the late currents.

Using the pulse methdology, we were able to generate four light-response curves: two using light delivered from the microscope, and based upon the peak current (Fig 2B, black trace), and the “steady-state” current (mean of the final 100ms; Fig 2B, red trace)) respectively, and two equivalent curves from illumination by the LED (Fig 2D), which additionally passed through different thickness blocks of mouse brain tissue (Fig 2E,F). With the mean currents across the population of recordings, the relative difference between the microscope and LED illuminations (which was the key measure) were virtually identical for the calculations from the peak and steady state currents, but the variance was much less for the peak measures, so further analyses focused upon those. We made detailed analyses of 16 recordings, in which we were able to map out the entire light-response curves, extending well beyond saturation. We were thus able to normalise these light responses according to the saturating current. Reflecting the fact that the light-sensitive component, the ChR2 molecule was identical in all cases, these light-response curves were extremely reproducible, with very low variance between recordings (Fig 2E,F).

These measurements generated, for each nucleated patch, a pair of light response curves, one generated from the microscope, and a second from the LED and for which light additionally passed through different lengths of brain tissue. The abscissa values plotted for the LED were the values of the light intensity at the LED itself, at its source for specifed currents passing through the LED (1-5mA currents, =∼1-5.3 mW light output, estimated using an integrating sphere, Fig 1F; see Methods). Light then was dissipated by the experimental arrangement as it passed through several interfaces (air/glass/saline/tissue, all of which will contribute to the reduced illumination of the nucleated patch). This meant that the LED data was always shifted by some amount to the right of the microscope data, which was the true illumination level onto the nucleated patch. The correction therefore provides a measure of light dispersion between the LED and the nucleated patch.

We derived a best fit from the microscope data. Importantly, simply shifting the LED data always created an excellent alignment with the microscope data (Fig 3A). We calculated the “correction” to achieve an optimal match for each pair of light-response curves, to provide an estimate of the effective light attenuation from the LED light source. This correction, in terms of the log-units of illumination, was then plotted with respect to the thickness of the brain tissue (Fig 3B). Significant attenuation of the light signal was observed in the absence of neural tissue in the bath.

**Figure 3.**
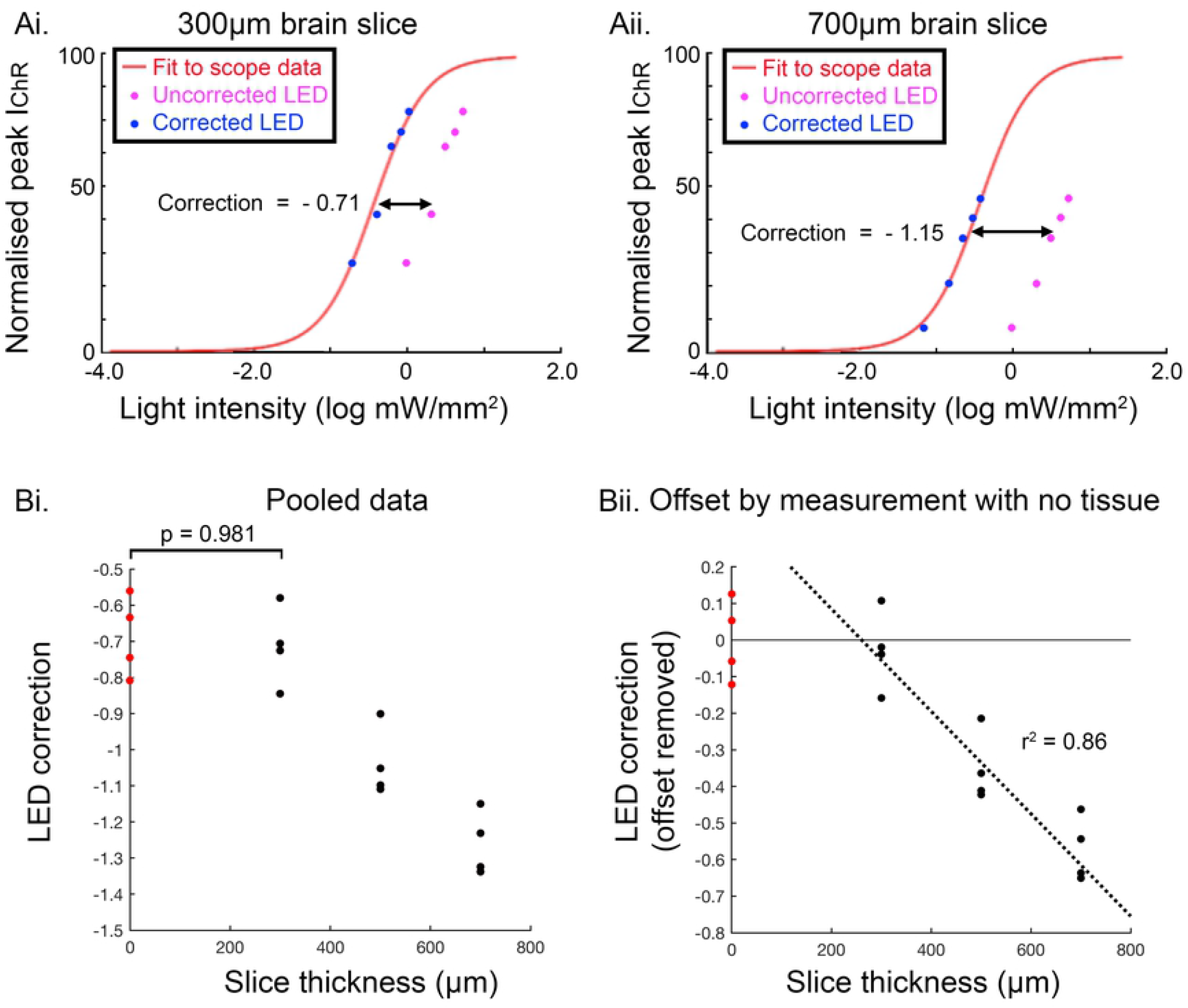
Aligning LED illumination and microscope illumination to estimate light penetration through cortical tissue. (A) Illustration of the derivation of the “LED correction” applied to data sets with light penetration through either (Ai) 300µm brain tissue or (Aii) 700µm brain tissue. The fit from the “ground-truth” microscope measurements, for which we have precise illumination intensity at the location of the nucleated patch (the focal plane of the microscope objective) is shown in red. The correction is how far this best fit needs to be shifted, to provide the best fit for the LED data (for which illumination measurements are imprecise). (Bi) Pooled data for all nucleated patches (16 cells, 4 measures taken without brain tissue, and 4 each for brain slices of 300µm, 500µm and 700µm thickness). (Bii) The same data set, offset by the mean of the data taken without a brain slice, to correct for other experimental sources of reduced light delivery (light dispersion, passing through the bottom of the recording chamber). We made a linear fit (note, however, that the ordinate scale is logarithmic), which has a gradient of -1.4 log_10_ units/mm of tissue (1 log unit drop in 735µm), indicating a space constant, *τ* = 319µm.

Assessing the whole data set, there was a highly significant effect of brain slice thickness (1-way ANOVA, F = 29.49, p<0.001), with significantly larger corrections required for increasing slice thickness, indicative of progressive attenuation of the light beam as it passes through the tissue. Pairwise comparisons for each group showed highly significant differences for every comparison (Table 1), except between the data for 300µm brain slices and measures made without a brain slice (p = 0.981). It is relevant that the way the recordings were made – nucleated patches were pulled from cultured neurons on a glass coverslip, and while they were then moved directly above the LED, they were not moved closer to it in the z-axis (see discussion, and Fig 5A) - the actual distance of the nucleated patch from the LED, in the recordings without a brain slice, was similar to those with a 300µm brain slice.

**Table 1.** Comparison of the mean correction calculated from light passing through brain slices of varying thickness. All p-values were calculated with a Tukey-Kramer test.

| Pairwise Comparisons | p-value |
| --- | --- |
| 0 µm vs 300 µm | 0.981 |
| 0 µm vs 500 µm | **0.002** |
| 0 µm vs 700 µm | **< 0.001** |
| 300 µm vs 500 µm | **0.003** |
| 300 µm vs 700 µm | **< 0.001** |
| 500 µm vs 700 µm | **0.041** |

**Table 2.**
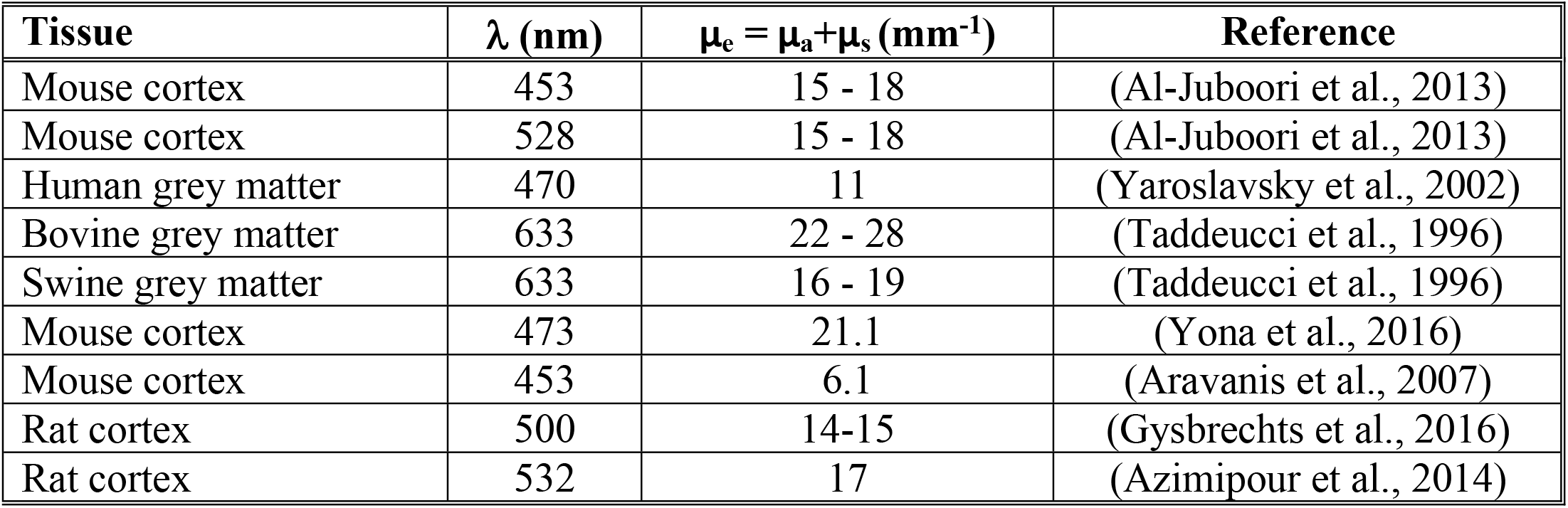
Extinction coefficient measurements from previous studies. Note that the extinction coefficient, µ_e_ is approximately equal to the scatter coefficient, µ_s_ because the absorption coefficient, µ_a_, is much smaller than µ_s_.

| Tissue | $\lambda$ (nm) | $\mu_e = \mu_a + \mu_s$ ($\text{mm}^{-1}$) | Reference |
| --- | --- | --- | --- |
| Mouse cortex | 453 | 15 - 18 | (Al-Juboori et al., 2013) |
| Mouse cortex | 528 | 15 - 18 | (Al-Juboori et al., 2013) |
| Human grey matter | 470 | 11 | (Yaroslavsky et al., 2002) |
| Bovine grey matter | 633 | 22 - 28 | (Taddeucci et al., 1996) |
| Swine grey matter | 633 | 16 - 19 | (Taddeucci et al., 1996) |
| Mouse cortex | 473 | 21.1 | (Yona et al., 2016) |
| Mouse cortex | 453 | 6.1 | (Aravanis et al., 2007) |
| Rat cortex | 500 | 14-15 | (Gysbrechts et al., 2016) |
| Rat cortex | 532 | 17 | (Azimipour et al., 2014) |

For all measurements through brain slices, the nucleated slice was located immediately above the brain slice. As such, both the physical distance from the LED, and the distance travelled through brain tissue increased by the same amount. These data showed that with increasing thickness of brain slices, there was a highly significant drop in the light current (Table 1), and collectively (excluding the “no brain slice” measurements since these were qualitatively different – change in distance travelled through brain, but no change in LED distance), this reduction in light current was linear, when plotted on a logarithmic scale (Fig 3B; gradient = - 1.37 logunits/mm; 95% confidence interval, -1.75 to -0.985). This corresponds to a 10-fold drop in illumination when light traverses 732µm of tissue, indicative of an exponential decay with a space constant = 317µm (95% CI, 248-441µm).

### Direct visualization of light attenuation passing through brain tissue

We compared the nucleated patch measurements to our previously published data derived by visualising the illumination directly (Dong et al., 2018). Our approach is illustrated in Fig 4. In short, we observed the pattern of scattered light, orthogonal to the direction of light penetration. Photomicrographs of the tissue allowed estimates of the illumination contours, as it was attenuated, with increasing distance from the source (Fig 4E). Measuring along the central line of illumination, we estimated the attenuation of light intensity to fall exponentially with a space constant of 453µm (95% confidence interval = 304-895µm, Fig 1F). This represents about a 36% increase over the nucleated patch measures, although note that the 95% CIs of both measures show considerable overlap.

**Figure 4.**
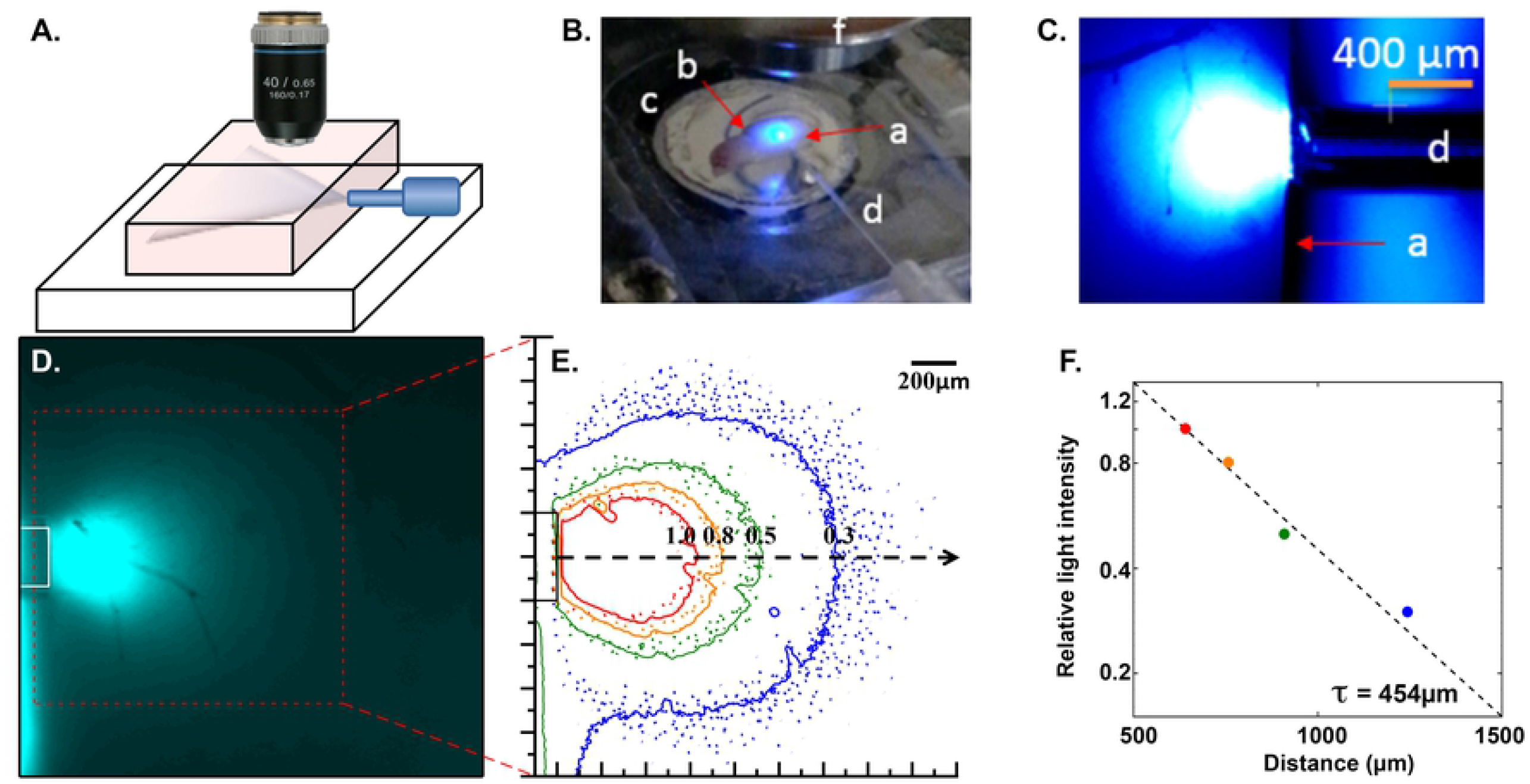
Estimating light penetration through tissue, by direct visualization. (A) Schematic illustration of the experimental arrangement, showing the illumination of a rodent brain slice from its lateral edge, and the observation of the light penetration from above. (B) Photograph of the experimental arrangement, showing the brain slice (a-b), located in a perfusion bath (c), with light being delivered through an optic fibre (d). (C and D) Visualization through the microscope objective. (E) Contour map of the visualised light spread through the tissue, viewed from above. (F) Relative light intensity (normalised to the red data points), orthogonal to the centre of the illumination (dotted line in E).

## Discussion

We have presented two different approaches to estimating light penetration through brain tissue, one utilizing nucleated patches, as biological light sensors, and the other involving direct visualization of the light, orthogonal to the light source. An added benefit of the former approach is that the measuring device is itself highly relevant to the optogenetic application, since it involves optogenetic proteins embedded inside the patch of membrane. This showed appreciable activation of the protein by LED illumination passing through hundreds of microns of brain tissue.

There are, however, some interpretative difficulties associated with both measures, as illustrated in the schematics in Fig 5, and which account for the small differences between the estimates of light attenuation derived from the two experimental approaches. The direct illumination estimate is compromised by significant reflection back into the tissue, from the cover slip underneath, which the light hits at a very shallow angle (Fig 5A). This serves to focus more light onto the distant tissue, than arrives directly, and consequently, the decay profile is probably steeper.

**Figure 5.**
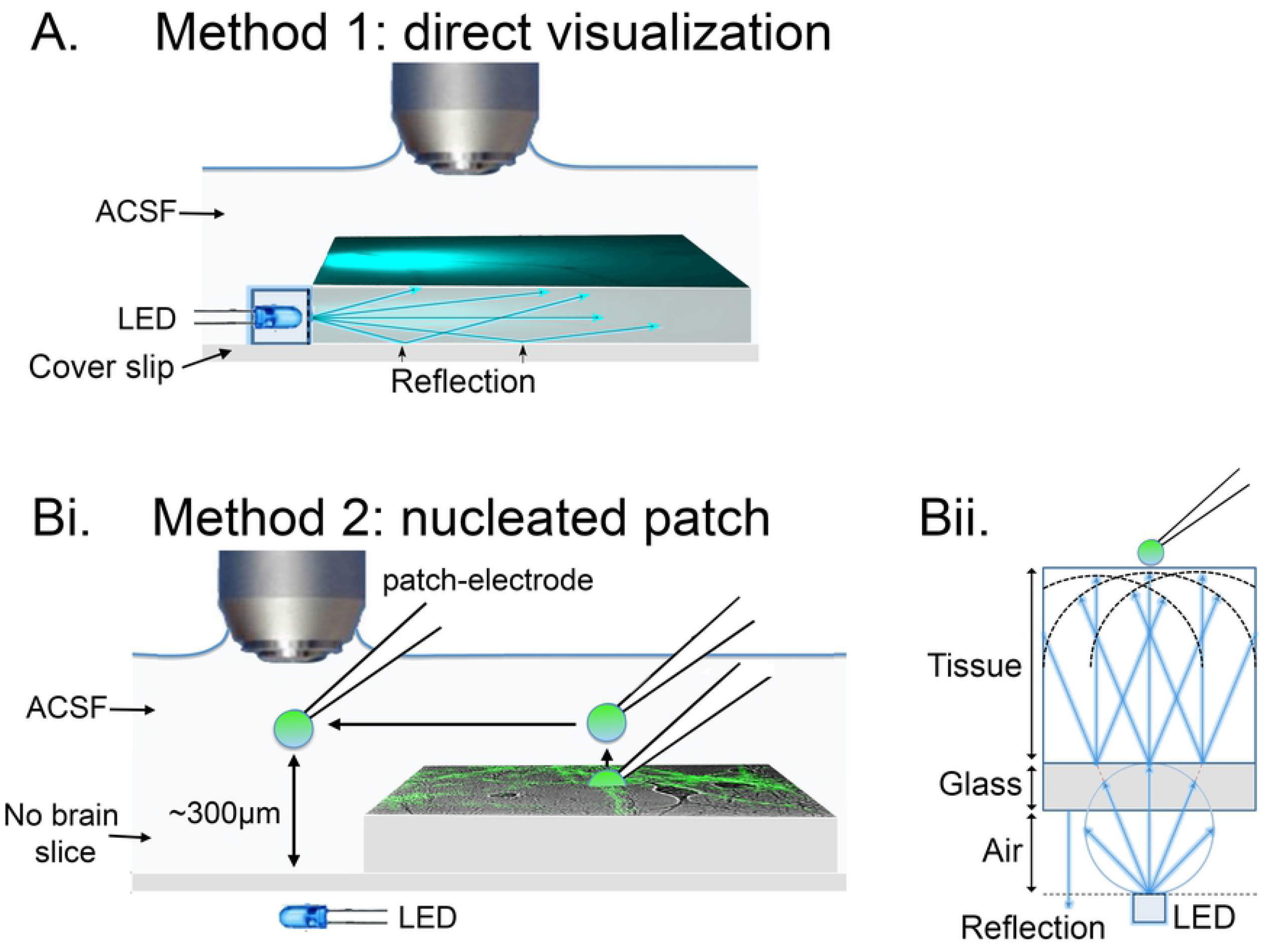
Factors affecting the delivery of light from LED. (A) Schematic illustrating the reflection of the bottom of the tissue chamber, that would lead to an overestimation of how far light spread through the tissue. (Bi) Schematic illustrating the nucleated patch experimental arrangement, when there was no brain tissue. The important detail is that, because the nucleated patches are pulled from cultures grown upon a glass slide, and then translocated only in a horizontal direction, the effective distance from the LED is almost exactly as for the case when a 300µm brain slice is present. Moving the nucleated patch closer to the light source is extremely problematic, because the glass bottom of the chamber is invisible, and electrodes are very easily broken on it. (Bii) Schematic illustrating the scatter of light. Note that the light will be scattered away from the direct illumination path to the nucleated patch, but this effect is offset to some considerable degree by light being scattered in a forward direction, back on to the patch. This light follows a longer path, of course, but the current measured does not distinguish this fact.

The decay in irradiance (i.e. radiant energy per unit area) is typically modelled as an exponential decay, as per the Beer-Lambert Law (Vo-Dinh, 2015):

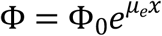

Where Φ represents the photon flux intensity. µ_e_ is the extinction coefficient and consists of both scattering and absorption i.e. *μ*_*e*_ = *μ*_*s*_ + *μ*_*a*_. If we assume an exponential relationship, then the space constant (τ = 1/extinction coefficient, µ_e_) is estimated to be 453µm (µ_e_ = 2.2mm^-1^), although as just mentioned, this may be an overestimate.

The nucleated patch experiments presented a different anomaly, because we measured no apparent difference in illumination intensity measurements with 300µm brain slices versus those without any intervening brain slice. This can be explained in two ways: first, the actual distance from the LED was the same in both cases, and second, the main scattering is forward, meaning that while light might be scattered away from the nucleated patch, other light is scattered on to it, meaning that the total amount of light was barely changed. For the 500µm, and 700µm brain slice recordings, the nucleated patch was further from the LED (in all cases, the nucleated was immediately above the slice), and so these did represent both an increased physical distance from the LED, and also further distance that light traversed brain tissue. In short, the measurements without brain slices clearly represent a different case, so we excluded these from further analyses, and calculated the effect of increasing brain slice thickness by fitting only the data sets with brain slices present, normalizing these to the 300µm brain slice measurements. These were linearly arranged, when plotted on a semi-logarithmic plot, indicating a space constant of 317µm (equivalent to an extinction coefficient, µ_e_ = 3.1mm^-1^).

Past exploration of the extinction coefficient in tissue (as per table 2) has shown that the scattering coefficient *μ*_*s*_ ∼ 10 – 30 mm-1, whereas the absorption coefficient *μ*_*a*_ ∼ 0.01mm-1. As such this component is typically ignored. There are some complications in that the tissue is not homogeneous. For example, Wang et. al. (Wang et al., 2017) has shown that the scattering coefficient varies significantly across the grey matter, but, more typically, scattering coefficients are taken as an average for tissue types – e.g. grey matter, white matter, for given animal species. Table 2 lists the recent measurements of µ_e_.

On initial inspection, our two approaches yield estimates of the extinction coefficient (µ_e_ = 2.2 and 3.1mm^-1^ respectively) that appear much smaller than these previous estimates (Table 2); that is to say, our data suggests that light penetrates further into the tissue, so it is important to consider what is the source of these differences.

A key consideration is the emission profile of the light, for a given light source. Notably, all the studies listed in Table 2 used a collimated light beam, as their light source, for which scattering causes divergence from the primary beam profile, thus diluting the irradiance (radiant power per unit area). To take this into account, Aravanis et al (2007) used a slightly different model, the Kubelka Munk model (Vo-Dinh, 2015) to explore light traversal.

In contrast, our study used an LED, which is not a collimated emitter; rather, it emits light in a Lambertian manner. In this case, light already emits at every angle with a spherical emission profile, as per Fig 5Bii, and this greatly reduces the effects of forward scattering. The reason for this is as follows. In homogeneous media, with particles much smaller than the wavelength of light, scattering will be isotropic; that is to say, all scattering directions have equal probability. This is known as Rayleigh scattering. However, in brain tissue, various particles including proteins and cellular organelles are larger than the light wavelength, resulting in “anisotropic scattering” with a significant preference for the forward direction. The directionality of the light scattering is given by the anisotropy term, g, which is a measure of the proportion of light still going forward, after a scattering event (Vo-Dinh, 2015). For the studies listed in Table 2, *g* is estimated to be 0.86 – 0.9 (Al-Juboori et al., 2013; Yaroslavsky et al., 2002; Yona et al., 2016). However, for LED emission profiles, the forward scattering does not significantly affect the beam decay, because almost as much light is scattered back into the direct path as is scattered out. The reverse backscattering thus becomes the dominant term, and can be estimated from g, by substituting the reduced scattering factor, 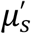 in place of *μ*_*S*_ (Hamdy et al., 2017), as follows:

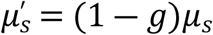

If we assume our measurements are primarily of backscattering, with relatively little effect from forward scattering, then to align our measurements to these previous ones, we should multiply our estimates of µ_e_ (2.2 and 3.1mm^-1^ respectively) by 1/(1-g) (i.e. by a factor of 7.1-10), which brings our estimates closely in line with those shown in Table 2. The notable conclusion, however, is that if one starts with a dispersed light source, the subsequent extinction profile appears to be far less steep than for a collimated light source.

One additional factor relevant to the *in vivo* situation, but not measured here, is the absorption of light by blood (Dunn, 2014). Given the highly anisotropic nature of the brain vascular supply at the sub-millimeter scale relevant to this discussion (Blinder et al., 2013), a generalized estimate of this effect is not easily measured empirically. Rather, the approach should be to model the length of the light path travelled, before encountering a blood vessel, for different tissues. This will further reduce the light penetration relative to our measurements, and this effect may be worsened if placement of an optrode induces local vascular reorganization.

## Acknowledgements

The project CANDO (Controlling Abnormal Network Dynamics with Optogenetics) is co-funded by the Wellcome Trust (102037) and Engineering and Physical Sciences Research Council (A000026). We thank all members of the CANDO consortium (www.cando.ac.uk).

